# An *in vitro* microfluidic culture device for peripheral neurotoxicity prediction at low concentration based on deep learning

**DOI:** 10.1101/2022.11.09.515778

**Authors:** Xiaobo Han, Naoki Matsuda, Kazuki Matsuda, Makoto Yamanaka, Ikuro Suzuki

## Abstract

In this study, a microfluidic culture device was developed including related evaluation methods using deep learning for the purpose of constructing a rapid assessment platform for peripheral neuropathy caused by compounds. Primary rodent dorsal root ganglion was cultured in the microfluidic culture device that could separate the cell body and neurites, and the neurites’ morphological changes were analyzed by immunofluorescence images. Separated neurites successful culture in the microfluidic device for more than 1 month indicated that a series of test processes from culture to drug stimulation and fluorescence observation is possible. Additionally, cultured samples were treated with several anticancer drugs known to cause peripheral neurotoxicity (i.e., vincristine, oxaliplatin, and paclitaxel) and analyzed the neurites’ morphological changes by deep learning for image analysis. After training, artificial intelligence (AI) could identify neurite morphological changes caused by each compound and precisely predicted the toxicity positivity, even at low concentrations. For testing compounds, AI could also precisely detect toxicity negative and positive based on neurite images, even at low concentrations. Therefore, this microfluidic culture system is supposed to be useful for *in vitro* toxicity assessment.

## 1. Introduction

Chemotherapy-induced peripheral neuropathy (CIPN) is one of the major common adverse chemotherapy events that is primarily associated with neurological abnormalities linked to pain, loss of sensation, and motor functionality, which ultimately leads to a decreased quality of life [1-3]. CIPN diagnosis and treatment remains as a challenging problem because its clinical presentation and molecular mechanisms are heterogeneous, and international guidelines has no clear consensus thus far [4,5]. An accurate assessment is thought to be essential to improve knowledge around CIPN incidence. Recently, there has been an increase in the number of *in vitro* cell models of rat and mouse dorsal root ganglia (DRG) sensory neurons to study CIPN at a mechanistic level [6-8]. Drug-induced neurotoxicity to DRG neurons under most commonly used chemotherapeutic agents (e.g., taxanes, vinca alkaloids, and bortezomib) was evaluated based on cell viability analysis and morphology [9]. For example, both bortezomib and vincristine-treated neurons reportedly showed decreased neurite outgrowth without increased cell death [10], while cisplatin and oxaliplatin treatment induced cell death [11]. Therefore, cultured DRG neurons can provide a reliable and powerful *in vitro* model for mechanistic and therapeutic CIPN studies.

However, traditional neurite outgrowth or cell viability assays are mainly efficient for drug dosing during or after CIPN symptom appearance. It is nearly impossible to conclude slightly morphological responses from neurites based on traditional assays, whereas such information could be important for CIPN prediction and prevention [9]. With deep learning technology development, artificial intelligence (AI) is enabling faster and easier information extraction from images [12]. In this study, an assessment platform for DRG neurotoxicity prediction was attempted to be established based on deep learning for image analysis. Here, a cyclo olefin polymer (COP)-based microfluidic device was used for DRG culture providing advantages, such as stable microstructures, chemical compounds absorption reduction, and mass production suitability [13]. Deep learning for image analysis was used to evaluate morphological changes in neurites images exposed to chemotherapeutic agents known to cause peripheral neuropathy, and its potential for drug-induced nerve abnormalities prediction was demonstrated even at a low concentration.

## Material and methods

### Device fabrication

Ushio Inc. prepared the microfluidic device as previously described [13]. Briefly, the vacuum ultraviolet (VUV) photobonding from an excimer light at a 172-nm wavelength was used to generate the functional groups (i.e., hydroxy and carboxyl groups) for assembling two COP material layers directly with heat treatment. One microfluidic device is comprised of four individual microfluidic cell culture channels (Figure 1A-a). The middle narrow slot part of the channel is 1000 µm in width, 165 µm in length, and 40 µm in height, with an open-top channel (1000 µm in width, 6 mm in length) and two circular holes (2 mm in diameter) at both ends which open to a rectangular medium reservoir (15 mm in width, 8 mm in length, and 5 mm in height). The maximum volume in each channel containing reservoirs is 1 mL. COP material (Zeonex 690R, Zeon, Tokyo, JAPAN) was injected to the two molds individually. The components were irradiated with VUV from an excimer lamp (172 nm; Ushio Inc., Tokyo, JAPAN) at 25°C after taking the structured COP components from the molds. The component surfaces were assembled using a heat press at less than 132°C. Finally, ethylene oxide gas (Japan Gas Co. Ltd., Kanagawa, JAPAN) was used for device decontamination.

**Figure 1.**
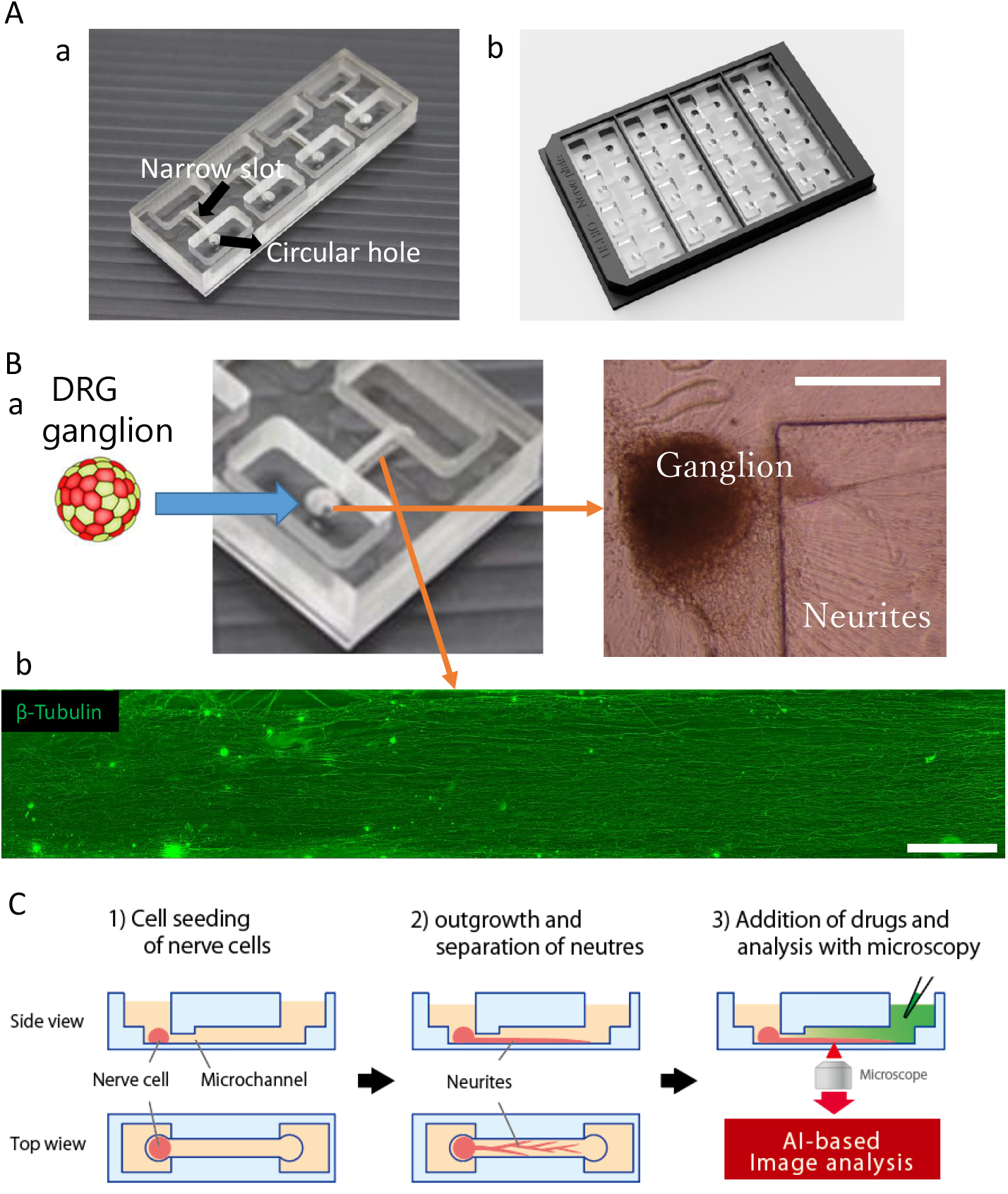
COP-based microfluidic device application on cell culture. A) Microfluidic device overview. B) A representative image of DRG ganglion cultured in the device. a) Phase contrast image of DRG ganglion in the seeding hole after 6 weeks of culture. Scale bar = 500 μm. b) Whole length immunofluorescence image sample of neurite outgrowth in the microfluidic channel after 6 weeks of culture. Scale bar = 500 μm. C) A schematic image of the experimental processes from cell culture to immunofluorescence observation.

### Cell culture

Before cell seeding, the microfluidic device was coated overnight at 4°C with 0.02% Poly-L-lysine (P4707, Sigma-Aldrich). The device was coated with 2.5 µg/mL laminin 511 (381-07363, Wako) for 30 min at 37°C after washing with phosphate-buffered saline (PBS).

DRG neurons were harvested as described previously [11]. The ethical approval for this study was obtained from Tohoku Institute of Technology Animal Care and User Committee. Briefly, timed pregnant female Wistar rats were anesthetized with isoflurane and then decapitated. E15 rat pups were removed from the uterus. Spinal ganglia were carefully removed, then immediately seeded into the circular hole at one end of the channel (one ganglion/channel). Cultures were incubated for the first 2 weeks in the Neurobasal Medium (21103049, Gibco) containing B27 supplement (17504044, Gibco), supplemented with 100 ng/mL NGF (01–125, Millipore). The medium was then replaced with the DMEM/F12 medium (11320033, Gibco), supplemented with 15% FBS, 50 µg/mL L-ascorbic acid, and 50 ng/mL NGF. This protocol yielded a stable neurites outgrowth population into the microchannel.

After another 6 weeks, three typical anticancer drugs were administrated to the cultures at two different concentrations for each: paclitaxel at 1 µM (paclitaxel) and 100 nM (paclitaxel low); vincristine at 30 nM (vincristine) and 3 nM (vincristine low); oxaliplatin at 100 µM (oxaliplatin) and 10 µM (oxaliplatin low). Acetaminophen at 10 µM, and sucrose at 100 µM as two negative drugs and DMSO at 0.1% as a control drug were also added to the cultures. Drug exposure lasted for 24h at 37°C.

### Immunocytochemistry

Sample cultures were fixed with 4% paraformaldehyde in PBS on ice (4°C) for 10 min. Fixed cells were incubated with 0.2% Triton-X-100 in PBS for 5 min, then with preblock buffer (0.05% Triton-X and 5% FBS in PBS) at 4°C for 1 h, and finally with preblock buffer containing a specific primary antibody, mouse anti-β-tubulin III (1:1000, T8578, Sigma– Aldrich) at 4°C for 24 h. Samples were then incubated with the secondary antibody, anti-mouse 488 Alexa Fluor (1:1000 in preblock buffer, ab150113, Abcam) for 1 h at room temperature. Stained cultures were washed twice with preblock buffer and rinsed twice with PBS. A confocal microscope (Eclipse Ti2-U, Nikon) for local images and an All-in-One fluorescence microscope (BZ-X, Kenyence) for whole microchannel-length images were the two microscope types used to visualize immunolabeling. The ImageJ software (NIH) was used to adjust the image intensity.

### Deep learning for image analysis and toxicity positive prediction

First, whole microchannel-length immunofluorescence images were subdivided into 50 × 50 pixel images and only the images in which axons were reflected were extracted to create the axon image dataset (Figure 3A). The training image dataset for AI analysis consisted of DMSO (n = 3 well), acetaminophen (n = 3 well), oxaliplatin (n = 4 well), and vincristine (n = 4 well). The untrained images dataset consisted of sucrose (n = 3 well), oxaliplatin low (n = 3 well), vincristine low (n = 3 well), paclitaxel (n = 3 well), and paclitaxel low (n = 3 well).

An AI for peripheral neurotoxicity prediction was created by transfer learning an image recognition model, GoogLeNet. AI was trained with DMSO and acetaminophen as negative control compounds and oxaliplatin and vincristine as positive control compounds (Figure 3A). The Leave-One-Out method was used to evaluate the classification algorithm performance. Only one well was excluded from the training dataset as validation data, used the trained images from the other wells to create an AI, and created an AI for each excluded well.

The training compounds’ prediction accuracy was calculated by averaging the unlearned wells’ prediction probabilities that were excluded from the training. Prediction accuracy for unlearned compounds was calculated by averaging the prediction results of multiple created models. The positive probability of a single well was calculated from the percentage of positive results out of the number of segmented images.

### Statistical analysis

The one-way ANOVA was used to perform multiple group comparisons followed by Dunnett’s test or Holm’s test, which were used to calculate the significant difference between each concentration.

## Results

### Application of the microfluidic device in cell culture

The microfluidic device was created by direct COP materials bonding, providing high mechanical stability and probability for a scaled paralleling evaluation platform (Figure 1A-b). DRG ganglion was seeded into the circular hole at one end of the microfluidic channel, and no cell damage or detachment was observed during culture (Figure 1B-a). After immunostaining, a clear neurite image could be achieved for the whole microchannel-length (Figure 1B-b). Neurites were confirmed to have sufficiently grown to occupy almost the whole microfluidic channel arear and the axon elongated unidirectionally along the horizontal direction. Therefore, the COP-based microfluidic device provides a possibility to perform test processes from cell culture to immunofluorescence observation (Figure 1C).

### DRG morphological presentation

Figure 2 shows the local immunofluorescence images of cultured DRG neurons at 8 weeks in the microfluidic channels after drug administrations. After vincristine administration, morphological abnormalities could be clearly confirmed including axon decrease and fragments (Figure 2B-a). However, such morphological change was not observed under vincristine low (Figure 2B-b). Also, after paclitaxel and oxaliplatin administration (Figure 2C, 2D), there was no significant morphological change in the neurite images. Since the neurite density remained at quite a high level under all conditions, a large datasets scale could be achieved enough for the following AI analysis.

**Figure 2.**
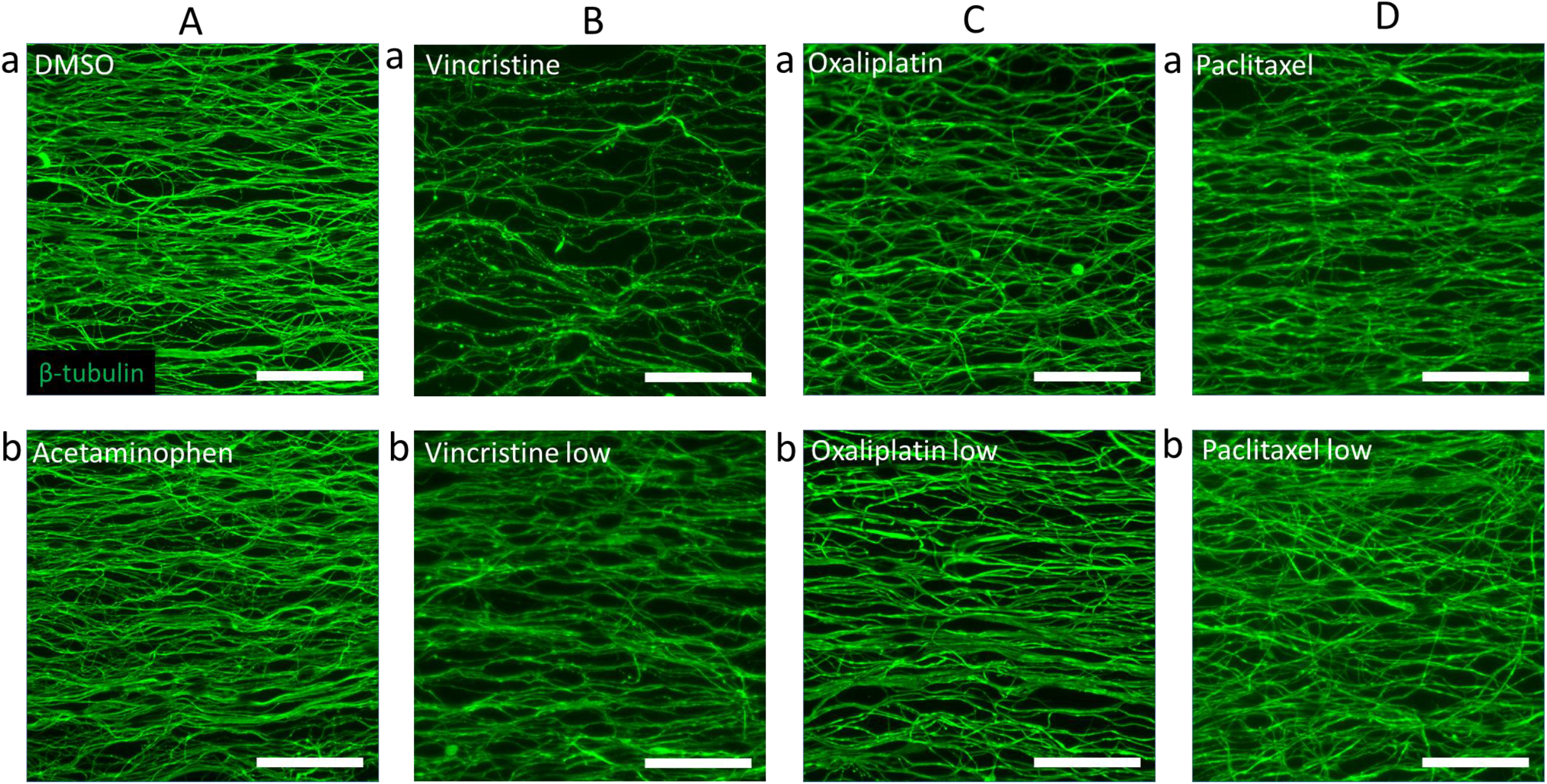
Representative local immunofluorescence image samples of neurites in the microfluidic channel after drug administration. Scale bar = 50 μm. A) a) 0.1% DMSO (control) and b) 10 µM acetaminophen (negative). B) a) 30 nM vincristine and b) 3 nM vincristine (vincristine low). C) a) 100 µM oxaliplatin and b) 10 µM oxaliplatin (oxaliplatin low). D) a) 1 µM paclitaxel and 100 nM paclitaxel (paclitaxel low).

### Neurotoxicity prediction using deep learning for images analysis

An AI for peripheral neurotoxicity prediction was created and trained with neurites image datasets as described above (Figure 3A and B). Figure 3C showed the prediction rate of toxicity positivity for each compound in training and testing groups. In the training group, both the control compound, DMSO (17.8%) and the negative compound, acetaminophen (13.5%) showed a low toxicity positive rate. A predicted positive line was set based on these two compounds, which is the average value of their positive rate plus 2 times of the standard deviation value that equaled to 42.4% (the red line in Figure 3C). As a result, both two positive compounds (i.e., oxaliplatin and vincristine) were toxicity positive detected even at a low concentration. The prediction accuracy was then tested with two unlearned compounds in the testing group. Sucrose was detected as a toxicity negative drug, while paclitaxel low was detected for toxicity positive significantly.

**Figure 3.**
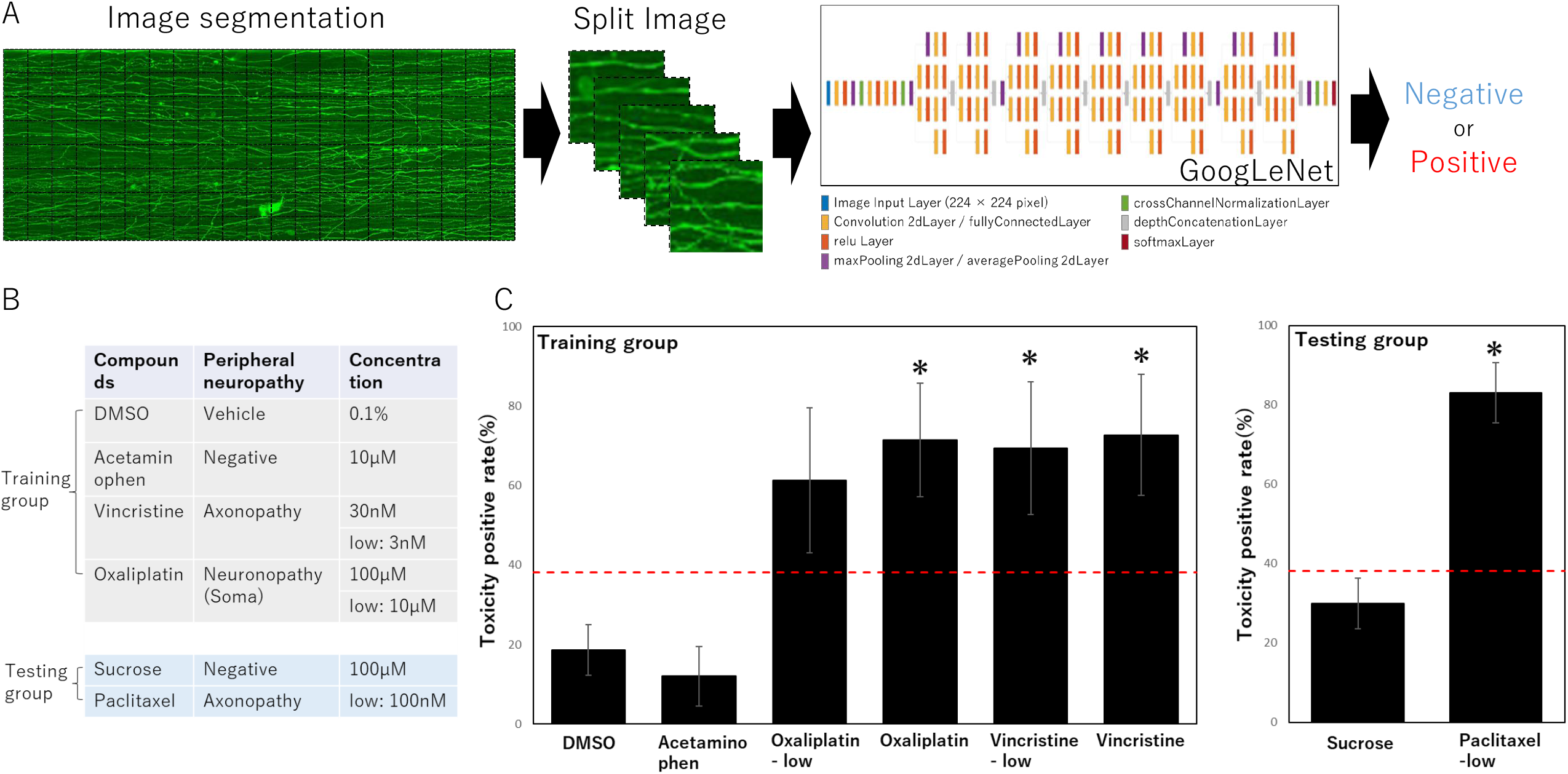
Neurotoxicity-positive AI prediction based on deep learning for neurite images analysis. A) Dataset treatment for deep learning and the AI analysis process. B) Detailed list of drug administration in the present study. C) The neurotoxicity positive rate predicted by AI analysis for each drug. Data were expressed by means ∓ standard errors. A toxicity positive line (red line in the graph) is determined as the average positive rate of two negative control compounds (i.e., DMSO and acetaminophem) plus two times of standard deviation, which is 42.4%. ANOVA was used to perform statistical analysis followed by post hoc Dunnet’s test or Holm’s test, *; p < 0.05 vs. negative.

## Discussion

*In vitro* cell models have been commonly used to explore presenting symptoms and underlying mechanisms of CIPN [8,9]. The recent development of biomaterial-based microphysiological systems provide efficient platform for drug screening and neurotoxicity assessment for drug candidates, as an alternative to animal models [14-16]. Of these materials, the COP-based device has several advantages such as chemical/physical stability and high optical clarity [17]. The COP-based device has been proven to maintain normal stem cell growth without undesired cellular damage [13]. In this study, a COP-based microfluidic device for primary rodent DRG cultures was developed demonstrating its potential as an *in vitro* peripheral neurotoxicity assessment platform with an AI analysis method using neurite images.

Primary rodent DRG neurons are commonly used to study CIPN induced by platinum derivatives, vinca alkaloids, and toxoids [6-11]. The DRG culture model is a heterogeneous system containing multiple cell types residing in a dorsal root ganglion, and therefore represents a more physiologically relevant model than those using highly purified neuronal cell lines or human iPSC-derived neurons. However, this feature also brings difficulties for evaluating the effects of agents using one simple standard. In the present study, the microfluidic device allowed only neurite elongation into the narrow channel. Therefore, we could only focus on the morphological changes in neurites, which are thought to be more susceptible than neuronal cell bodies to agents that cause CIPN.

Vincristine at a 30-40 nM concentration could reportedly induce axonal degeneration in cultured primary DRG neurons after a 24-h exposure [8,10], which is comparable with our results (Figure 2B-a). In the other hand, oxaliplatin administration over 33.2µM could reportedly significantly decrease neurite outgrowth in DRG explants [11], which was not observed in the present study. Indeed, different compounds could proceed via distinct mechanisms, as the cell body is accordingly required for oxaliplatin-induced axon degeneration [18,19]. Therefore, reproduction of similar results was challenging using the present microfluidic channel, in which the neurites were separated from the cell body. However, after deep learning for image analysis, oxaliplatin toxicity could still be predicted using neurite images at a relatively high accuracy.

One notable advantage of AI image analysis is that a high accuracy for toxicity positive prediction of vincristine and oxaliplatin was demonstrated at a relatively low concentration. In the previous researches, DRG cultures were detected as no significant changes or not reported under similar conditions [8,9]. However, the detailed characteristics of slight morphological changes could be reflected by deep learning for image analysis. This finding might lead to a new understanding of CIPN’s underlying mechanisms, and could be helpful for CIPN preventing at an early stage.

The prediction accuracy of our AI analysis method was also tested with unlearned compounds including paclitaxel, a widely used anticancer drug known to induce CIPN. Paclitaxel’s most toxic dose is 1 µM as previously reported [20]. The neurotoxic effect of paclitaxel is demonstrated mainly in DRG explants with necrosis occurrence. Neurite outgrowth reportedly has no significant changes after a 24-h paclitaxel exposure at a concentration range from 10 nM–1 µM [21], which is similar to our current results (Figure 2D). Using AI image analysis, paclitaxel toxicity was successfully detected based on neurite images, even at a low concentration. Taken together, it is believed that the COP-based microfluidic device combined with AI image analysis should be effective as an *in vitro* toxicity assessment platform of peripheral neuropathies.

## Acknowledgement

This study was supported by the grant of Ushio Inc. as a collaborative project.

## References

[1] G. Cavaletti, P. Alberti, B. Frigeni, et al. Chemotherapy-induced neuropathy. Curr. Treat. Options Neurol. 13 (2011) 180–190. doi: 10.1007/s11940-010-0108-3.

[2] S. Eldridge, L. Guo, J. Hamre. A Comparative Review of Chemotherapy-Induced Peripheral Neuropathy (CIPN) in In Vivo and In Vitro Models. Toxicol. Pathol. 48 (2020), 190–201. doi: 10.1177/0192623319861937.

[3] H. Starobova, I. Vetter. Pathophysiology of Chemotherapy-Induced Peripheral Neuropathy. Front. Mol. Neurosci. 10 (2017), 174. doi: 10.3389/fnmol.2017.00174.

[4] L. A. Colvin. Chemotherapy-induced peripheral neuropathy (CIPN): Where are we now? Pain 160 (2019), S1–S10. doi: 10.1097/j.pain.0000000000001540.

[5] C. L. Loprinzi, C. Lacchetti, J. Bleeker, et al. Prevention and Management of Chemotherapy-Induced Peripheral Neuropathy in Survivors of Adult Cancers: ASCO Guideline Update. J Clin Oncol 38 (2020), 3325–3348. doi: 10.1200/JCO.20.01399.

[6] B. Malgrange, P. Delrée, J. M. Rigo, et al. Image analysis of neuritic regeneration by adult rat dorsal root ganglion neurons in culture: quantification of the neurotoxicity of anticancer agents and of its prevention by nerve growth factor or basic fibroblast growth factor but not brain-derived neurotrophic factor or neurotrophin-3. J Neurosci Methods 53 (1994), 111–122. doi: 10.1016/0165-0270(94)90151-1.

[7] I. H. Yang, R. Siddique, S. Hosmane, et al. Compartmentalized microfluidic culture platform to study mechanism of paclitaxel-induced axonal degeneration. Exp Neurol 218 (2009),124–128. doi: 10.1016/j.expneurol.2009.04.017.

[8] L. Guo, J. Hamre, S. Eldridge, et al. Multiparametric Image Analysis of Rat Dorsal Root Ganglion Cultures to Evaluate Peripheral Neuropathy-Inducing Chemotherapeutics. Toxicol Sci 156 (2017), 275–288. doi: 10.1093/toxsci/kfw254.

[9] Y. Fukuda, Y. Li, R. A. Segal. A Mechanistic Understanding of Axon Degeneration in Chemotherapy-Induced Peripheral Neuropathy. Front Neurosci 11 (2017), 481. doi: 10.3389/fnins.2017.00481.

[10] S. Geisler, R. A. Doan, G. C. Cheng, et al. Vincristine and bortezomib use distinct upstream mechanisms to activate a common SARM1-dependent axon degeneration program. JCI Insight 4 (2019), e129920. doi: 10.1172/jci.insight.129920.

[11] L. E. Ta, L. Espeset, J. Podratz, A. J. Windebank. Neurotoxicity of oxaliplatin and cisplatin for dorsal root ganglion neurons correlates with platinum-DNA binding. Neurotoxicology 27 (2006), 992–1002. doi: 10.1016/j.neuro.2006.04.010.

[12] S. Ravindran. Five ways deep learning has transformed image analysis. Nature 609 (2022), 864–866. doi: 10.1038/d41586-022-02964-6.

[13] M. Yamanaka, X. Wen, S. Imamura, et al. Cyclo olefin polymer-based solvent-free mass-productive microphysiological systems. Biomed Mater 16 (2021). doi: 10.1088/1748-605X/abe660.

[14] K. Wang, K. Man, J. Liu, et al. Microphysiological Systems: Design, Fabrication, and Applications. ACS Biomater Sci Eng 6 (2020), 3231–3257. doi: 10.1021/acsbiomaterials.9b01667.

[15] M. Campisi, S. E. Shelton, M. Chen, et al. Engineered Microphysiological Systems for Testing Effectiveness of Cell-Based Cancer Immunotherapies. Cancers (Basel) 14 (2022), 3561. doi: 10.3390/cancers14153561.

[16] M. Virumbrales-Muñozabc, J. M. Ayusoab. From microfluidics to microphysiological systems: Past, present, and future. Organs-on-a-Chip 4 (2022), 100015. doi: 10.1016/j.ooc.2022.100015.

[17] P. M. Kristiansen, A. Karpik, J. Werder, et al. Thermoplastic Microfluidics. Methods Mol Biol 2373 (2022), 39–55. doi: 10.1007/978-1-0716-1693-2_3.

[18] M. J. Geden, M. Deshmukh. Axon degeneration: context defines distinct pathways. Curr Opin Neurobiol 39 (2016), 108–115. doi: 10.1016/j.conb.2016.05.002.

[19] D. J. Simon, J. Pitts, N. T. Hertz, et al. Axon Degeneration Gated by Retrograde Activation of Somatic Pro-apoptotic Signaling. Cell 164 (2016), 1031–1045. doi: 10.1016/j.cell.2016.01.032.

[20] G. Nicolini, R. Rigolio, A. Scuteri, et al. Effect of trans-resveratrol on signal transduction pathways involved in paclitaxel-induced apoptosis in human neuroblastoma SH-SY5Y cells. Neurochem Int 42 (2003), 419–429. doi: 10.1016/s0197-0186(02)00132-8.

[21] A. Scuteri, G. Nicolini, M. Miloso, et al. Paclitaxel toxicity in post-mitotic dorsal root ganglion (DRG) cells. Anticancer Res 26 (2006), 1065–1070.

